# Self-fertilization reverses the direction of selection on recombination

**DOI:** 10.64898/2026.08.18.745567

**Authors:** Tom Parée, Noah Sénéchal Chevalier, Denis Roze, Henrique Teotónio

## Abstract

The evolution of recombination is thought to be influenced by many factors, including the mating system. Here, we provide an experimental test of how self-fertilization (selfing) affects the evolution of a recombination modifier. We used experimental populations of *Caenorhabditis elegans* segregating for the recombination modifier *rec-1*, a mutant that redistributes crossovers from the genetically diverse chromosome arms toward the less diverse central regions. By evolving populations under varying selfing rates, we show that increasing selfing reverses selection acting on the *rec-1* mutant, from positive to negative. Simulations show that this reversal can be explained by an expansion of the genomic region over which the modifier remains associated with the genetic combinations it creates. These results demonstrate that selfing can fundamentally alter the evolutionary fate of recombination modifiers and reveal a mechanism not predicted by previous theoretical models of recombination evolution under different mating systems, which assumed uniform recombination landscapes.

## Introduction

Meiotic recombination rates, mainly determined by crossover frequencies, vary across species and individuals (Mercier et al., 2015; Stapley et al., 2017; Johnston, 2024). Crossover positions are non-uniformly distributed across the genome, creating hetero-geneous “recombination landscapes” (Haenel et al., 2018; Brazier and Glémin, 2022, 2024). Alleles modifying the number and position of crossovers underlie differences in recombination landscapes within and between species (Brand et al., 2018; Samuk et al., 2020; Johnston, 2024). Such recombination modifiers can be under direct selection because the number and distribution of crossovers affect meiosis and fertility (Hassold and Hunt, 2001; Hunt, 2006). Recombination modifiers can also be indirectly selected through the more or less fit genetic combinations they generate (Otto, 2021). Two factors determine the fate of a recombination modifier under indirect selection.

The first factor is the fitness effect of the genetic combinations produced by recombination. By freeing beneficial alleles from deleterious alleles, recombination can increase fitness variance and reduce selective interference. Combinations of deleterious and beneficial alleles are expected to be common in most populations, although their frequency depends on population size, the number of selected loci (Otto and Barton, 2001; Iles et al., 2003; Barton and Otto, 2005; Roze and Barton, 2006; Roze, 2021), spatial heterogeneity, and epistasis (Barton, 1995; Lenormand and Otto, 2000). In addition to affecting the frequency of these genetic combinations in a population, epistasis also influences their fitness effects and, consequently, can offset the benefit of recombination (Barton, 1995; Neher and Shraiman, 2009).

The second factor determining the fate of a recombination modifier is the number of generations during which it remains associated with the genetic combinations it creates. This duration decreases rapidly with the genetic distance between the modifier and the generated genetic combinations (Otto and Barton, 1997; Roze, 2021; see Figure 1 in Parée and Teotónio, 2025 for an illustration). Consequently, the modifier rapidly loses association with most genetic combinations it creates across the genome, and only those generated in its close vicinity lead to substantial indirect selection. This small local interval may not be representative of the genome-wide effects of the modifier because recombination landscapes and genomic organization are heterogeneous. As a result, recombination landscape modifiers can be favored despite increasing genome-wide selective interference and impairing the rate of adaptation. We previously demonstrated this phenomenon using the *rec-1* mutant in genetically diverse, predominantly outcrossing domesticated populations of *Caenorhabditis elegans* (Parée et al., 2025).

**Fig 1.**
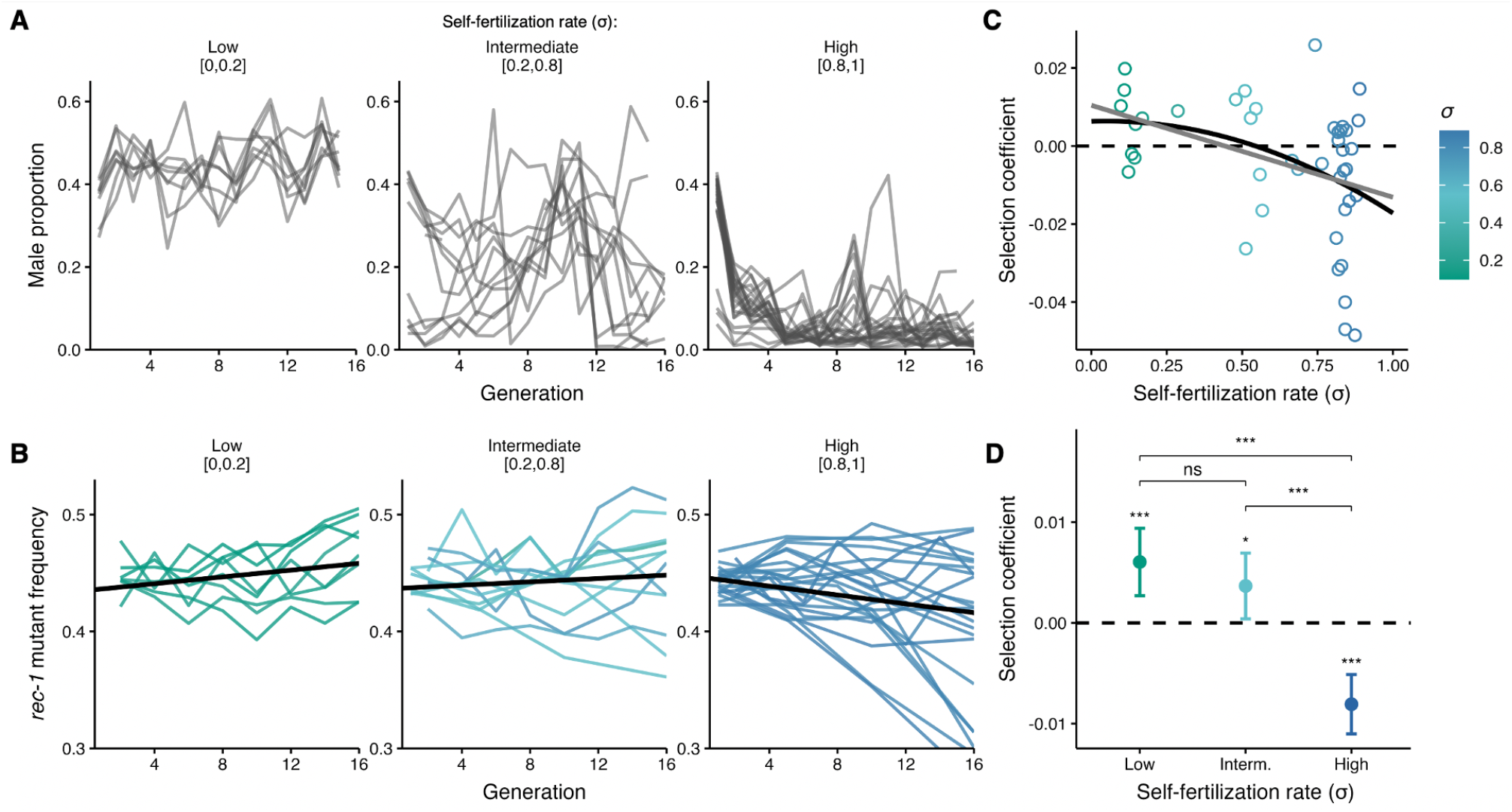
Experimental evolution. *A*. Male proportion during experimental evolution. Each line is an experimental population. Populations were classified into three arbitrary groups based on the arithmetic mean of their self-fertilization during evolution (low: n=8; intermediate: n=12; high: n=22). For a given sample, the self-fertilization rate is equal to one minus the male proportion in the next generation (Stewart and Phillips, 2002). *B. rec-1* allele frequency during experimental evolution. Each line is a replicate population. *C*. Each dot is the observed selection coefficient of the *rec-1* mutant for a given population. Lines represent the fitted regression line from the linear model with (black) and without (grey) a quadratic term. *D*. Dots and error bars are the estimated marginal mean and 95% confidence interval for each selfing rate bin estimated from a linear mixed model.

The *rec-1* mutant alters chromosomal crossover positioning without detectable direct fitness effects (Rose and Baillie, 1979; Rattray and Rose, 1988; Zetka and Rose, 1995; Parée et al., 2024, 2025). It increases genome-wide selective interference by redistributing crossovers from the genetically diverse chromosomal periphery toward the less diverse central regions, which exhibit low recombination rates in the wild type (Zetka and Rose, 1995; Parée et al., 2024). Despite reducing the rate of adaptation, the *rec-1* mutant is located within a genomic interval in which it locally increases recombination and is therefore favored because it reduces local selective interference (Parée et al., 2025).

Because the size of the local genomic interval driving indirect selection on a modifier depends on realized genetic distances, it may be affected by recombination itself, population structure, and mating system. Crossover frequencies positively correlate with self-fertilization (selfing) rates across flowering plant species (Roze and Lenormand, 2005; Ross-Ibarra, 2007; Brazier et al., 2025). In animals, however, this relationship has not been robustly tested because of the limited availability of recombination data in hermaphroditic species (Stapley et al., 2017), although this has not been updated recently. By increasing homozygosity, selfing reduces effective recombination and intensifies selective interference through stronger linkage disequilibrium, particularly in low-recombining genomic regions (Nordborg and Donnelly, 1997; Teterina et al., 2023; Burgarella et al., 2024; Lucek et al., 2025). Across a wide range of parameters, selection for recombination is predicted to be maximized at high selfing rates, and to vanish under complete selfing due to the absence of effective recombination (Stetsenko and Roze, 2022). However, these predictions have never been tested experimentally and were derived under the assumption of modifiers that alter recombination uniformly across a homogeneous genome. The dynamics may become substantially more complex when recombination landscapes are heterogeneous.

Here, we aim to better understand the impact of self-fertilization on indirect selection acting on a recombination modifier. Specifically, we extend our previous work using experimental *C. elegans* populations polymorphic at *rec-1*, which was conducted under predominantly outcrossing, by now incorporating varying selfing rates. We show that increasing the selfing rate not only alters the strength of selection acting on *rec-1* alleles, but also reverses its sign, leading to negative selection on the mutant allele. This reversal is mirrored in individual-based simulations and can be explained by the dramatic change in the span of the genomic region that drives indirect selection. Under predominant outcrossing, this span is restricted to a small interval surrounding *rec-1*, whereas under predominant selfing it extends across the entire genome because of the drastic reduction in effective recombination. Consequently, indirect selection acting on the mutant becomes more closely aligned with its genome-wide effect, which is to increase selective interference.

## Results

### Experimental evolution

We conducted evolution experiments to test for indirect selection on the *rec-1* mutant under varying selfing rates. In total, 42 genetically diverse experimental populations, polymorphic for *rec-1*, were maintained for 16 generations in a high-salt environment they had never encountered before. At specific time points during the experiment, selfing rates were manipulated by impairing male reproduction and enforcing hermaphrodite self-fertilization. This resulted in fluctuating male frequencies, with transient drops that typically lasted for a few generations before recovering and being controlled again (Figure 1A). The arithmetic mean of the selfing rate for each population was calculated and compared to the frequency trajectories of the *rec-1* mutant, measured at regular intervals (Figure 1B). Higher mean selfing rates affected the indirect selection of the *rec-1* mutant, reversing it from positive under predominant outcrossing to negative under predominant selfing (Generalized Linear Mixed Model: *p* − *value* = 3.1 ×0^*−*12^). Selection coefficients for the *rec-1* mutant, calculated for each population or different selfing rate bins, are presented in (Figure 1C,D). As expected, the variance in observed selection coefficients between populations increases with selfing rate (generalized least squares model: *p* − *value* = 2.9 × 10^*−*2^). The reversal of selection with increased selfing is robust when the analysis is repeated using the harmonic mean of the selfing rate instead of the arithmetic mean, giving more weight to smaller values (Figure S1).

### Simulations

To understand the mechanism behind the reversal of selection of the *rec-1* mutant under high selfing rate, we simulated the evolution of the *rec-1* mutant in androdioecious populations with varying selfing rates for 50 generations. Simulations mimic the experimental recombination landscapes as well as the haplotype diversity and SNV diversity observed in our experimental *C. elegans* populations (Figure 2A); i.e., lower diversity in the central regions where the *rec-1* mutant increases recombination rates. For computational efficiency, we simulated only three chromosomes: chromosome I (which harbors the *rec-1* gene), as well as chromosomes II and III. We observed that the selection on the *rec-1* mutant varies with the selfing rate. Under intermediate selfing (*σ* = 0.5), indirect selection on the modifier is increased compared to random mating, as expected due to the increased selective interference (Stetsenko and Roze, 2022). At a high selfing rate (*σ* = 0.9), the indirect selection of the *rec-1* mutant is reversed and becomes negative, echoing the experimental results.

**Fig 2.**
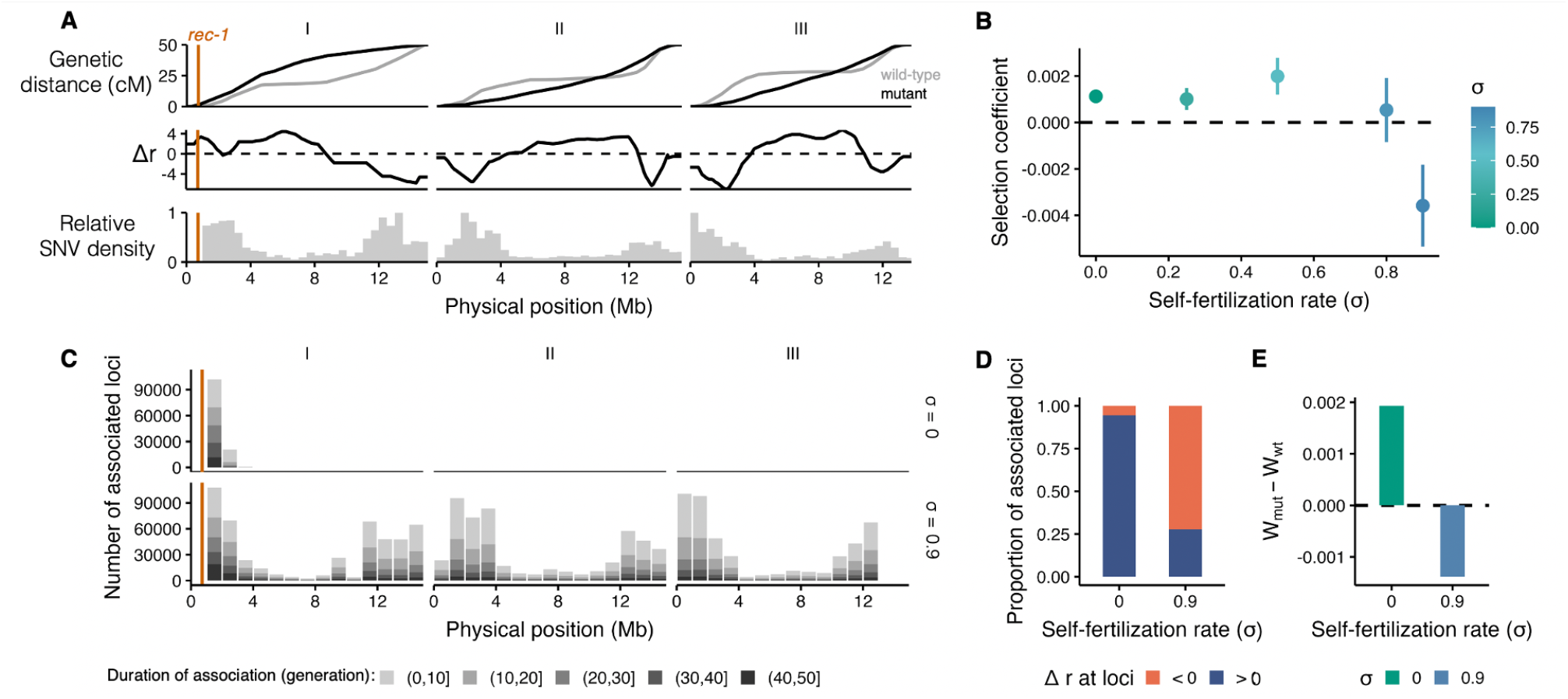
Simulations. A. The upper panel shows the *rec-1* wild-type (grey) and mutant (black) Marey maps from (Parée et al., 2024), where genetic distance is plotted as a function of the physical position of simulated chromosomes, and a greater slope indicates higher recombination rates. The middle panel shows the difference in recombination rate (*r*) between the mutant and the wild-type (Δ*r* = *r*_*mut*_ *− r*_*wild−type*_). The lower panel shows the relative SNV density for each simulated chromosome. The genomic position of the *rec-1* gene (I:719556) is indicated by the orange vertical line. **B**. Selection coefficient of the *rec-1* mutant at different self-fertilization rate across 2.3 *×* 10^4^ simulation runs with varying selfing rates. **C**. Number of loci building linkage with *rec-1* across all simulations in 1Mb genomic window in simulations with random mating (*σ* = 0) or high selfing rate (*σ* = 0.9). A more intense shade of grey indicates that the association lasted longer. At a given generation, a selected locus was determined to be significantly associated with *rec-1* when the absolute linkage disequilibrium was higher than a defined threshold (*D >* 0.033 or *D < −*0.033). This threshold corresponds to the maximum absolute value of *D* on chromosomes II and III under random mating, where linkage with *rec-1* is not expected. **D**. Proportions of associated loci that are found at a genomic position where the *rec-1* mutant increases recombination (Δ*r >* 0) or decreases it (Δ*r <* 0). **E**. Difference between the haplotypes associated with the *rec-1* mutant and wild-type during evolution under random mating or predominant selfing throughout evolution. For a given simulation run at a given generation, haploid genomes are classified depending on the *rec-1* allele they bear, mean fitness is computed, and the difference between the mutant and wild-type allele is calculated. This difference is then averaged across generations and simulations. A positive value indicates that, on average, the *rec-1* mutant allele is associated with fitter haplotype throughout evolution, and inversely.

Indirect selection on recombination modifier alleles arises when modifiers become associated with fitter (or less fit) genotypes because they alter the efficacy of selection. Hence, to understand the mechanism underlying the reversal of indirect selection on the *rec-1* mutant under high selfing rates, we quantified linkage disequilibrium (*D*) generated during evolution between the *rec-1* allele and selected loci across the genome under outcrossing (*σ* = 0) and predominant selfing (*σ* = 0.9). Under outcrossing, *rec-1* primarily establishes associations with nearby loci, as linkage with genotypes generated at distant loci decays rapidly (Figure 2C). Consequently, the associated loci sample only a narrow genomic interval in which the *rec-1* mutant predominantly increases recombination (Figure 2D). Under predominant selfing, reduced effective recombination dramatically expands the genomic scale over which *rec-1* becomes associated with selected loci, effectively all chromosomes. Associated loci are more frequently located in regions where the mutant decreases recombination because genetic diversity concentrates there (Figure 2C,D).

As a result, haplotypes associated with the *rec-1* mutant are, on average, fitter than those associated with the wild-type allele under outcrossing, but less fit under predominant selfing (Figure 2E). Indirect selection thus becomes negative when it integrates genome-wide effects, as it aligns with the overall effect of the *rec-1* mutant, which increases selective interference by redistributing crossovers away from the genetically diverse chromosomal periphery.

## Discussion

While direct selection likely constrains the diversity of meiotic recombination rates, there appear to be no mechanistic constraints on crossover formation that readily explain the elevated recombination rates observed in species compared to outcrossing ones (Ross-Ibarra, 2007). Instead, theoretical models have long predicted that self-fertilization should influence the indirect selection acting on recombination rates (Charlesworth et al., 1979). Here, we show that increasing selfing can not only alter the strength of indirect selection acting on a recombination modifier, but can also reverse its sign. In our system, this reversal occurs because the modifier’s effects on recombination vary across the genome, creating a misalignment between its local and genome-wide effects. Such misalignment may also arise whenever recombination is beneficial in some genomic regions but deleterious in others due to heterogeneity in genetic architecture across the genome.

Although selective interference appears sufficient to explain the main patterns observed here, pervasive epistasis has been documented in the *C. elegans* experimental populations used in this study (Chelo and Teotónio, 2013; Noble et al., 2017, 2021). Theory predicts that epistasis can strongly affect both selection on recombination modifiers and its relationship with selfing (Roze and Lenormand, 2005; Stetsenko and Roze, 2022). Co-adapted epistatic allele combinations may accumulate in ancestrally low-recombining regions (Neher and Shraiman, 2009; Venu et al., 2024), such as the chromosomal centers (see Parée et al., 2025). Other non-additive interactions, such as dominance, may also impact the evolution of *rec-1*. For instance, when deleterious alleles are recessive, low-recombining regions may enter a regime of pseudo-overdominance in outcrossing populations, but not in predominantly selfing ones (Sianta et al., 2023). Consequently, increasing recombination in chromosomal centers may have different consequences depending on the mating system. Untangling how these diverse genetic interactions shape selection on recombination modifiers remains an important challenge for future work.

Because indirect selection integrates effects across the entire genome under predominant selfing, it should become more closely aligned with the effect that the recombination modifier has on adaptation. In this sense, selfing may favor the evolution of more “optimal” recombination landscapes. The negative selection acting on the *rec-1* mutant under high selfing is consistent with our previous finding that the mutant impairs adaptation (Parée et al., 2025), although adaptive rates were only quantified in outcrossing populations. The shift from positive to negative selection on *rec-1* under predominant selfing may explain why *rec-1* loss-of-function alleles are absent from a large collection of wild *C. elegans* isolates (CaenDR; Crombie et al., 2024) despite lacking detectable direct fitness costs.

The framework presented here offers a new perspective on why selfing species often exhibit elevated recombination rates. In predominantly selfing populations, indirect selection should more strongly favor modifiers whose genome-wide effects enhance adaptation, which may often be those that increase genome-wide recombination. In contrast, the evolution of recombination in outcrossing species may be inherently more variable because selection acting on a modifier is largely contingent on the context within a relatively small genomic interval. Some intervals may experience strong selective interference, whereas others may lack genetic variation for fitness or contain co-adapted loci that disfavor recombination. The evolutionary stability of recombination landscapes in outcrossing species may also depend on whether recombination evolution is driven primarily by occasional sweeps of large-effect modifiers or by continuous selection acting on many small-effect modifiers whose combined local effects sample a much larger fraction of the genome.

The mating system should also influence the relative strength of indirect selection acting on cis-acting (local) versus trans-acting (genome-wide) recombination modifiers. In outcrossing species, indirect selection on both types of modifiers should be of a similar order of magnitude because it is driven primarily by their local effects anyway. In contrast, under predominant selfing, indirect selection on trans-acting modifiers integrates effects genome-wide and may therefore be substantially stronger than selection on cis-acting modifiers. This prediction is testable: mating systems should differ systematically in the proportion of recombination rate variation explained by cis- versus trans-acting modifiers. Whether such differences lead to more uniform recombination landscapes in self-fertilizing species remains an open question.

Nevertheless, difficulties arise when extrapolating from relatively short-term experimental evolution to long-term evolutionary dynamics in natural populations. Selfing species typically exhibit substantially reduced genetic diversity (Charlesworth, 2003; Glémin et al., 2006), which may ultimately weaken selection on recombination. In addition, mating systems are tightly intertwined with ecological conditions, demography, life-history traits, and genome architecture, complicating comparative analyses across species (Brazier et al., 2025). Within species, outcrossing rates can vary substantially (Teotónio et al., 2006; Felmy et al., 2023) and respond plastically to environmental conditions (Bishop et al., 2017), making them difficult to estimate accurately. Furthermore, outcrossing rates may themselves evolve under conditions that also favor increased recombination (Morran et al., 2009; Kamran-Disfani and Agrawal, 2014). One theoretical study suggested that selection acting on recombination modifiers is relatively weak compared to that acting on many other classes of modifiers (Proulx and Teotónio, 2022), although it did not consider modifiers of outcrossing rate. Because outcrossing and meiotic recombination both influence effective recombination, understanding their co-evolution will likely be essential for explaining recombination rate evolution in nature.

For decades, experimental tests of the extensive theoretical literature on recombination evolution have largely been limited to comparisons between sexual and asexual populations or to phenotypic measurements of recombination rates without knowledge of their underlying genetic basis (Parée and Teotónio, 2025). This study, together with our previous work on *rec-1* (Parée et al., 2025), represents the first evolution experiments to directly track a known meiotic recombination modifier. These experiments strengthen empirical support for indirect selection as a driver of recombination evolution and provide a powerful framework for testing longstanding theoretical predictions about conditions that shape the evolution of recombination.

## Methods

### Experimental evolution

To test indirect selection on a recombination modifier under varying selfing rates, we utilized the MR0 (Mixed Recombination G0) ancestral populations, which were previously generated (Parée et al., 2025). These populations were established through the mass introgression of wild-type and loss-of-function mutant alleles of *rec-1* into a genetically diverse, predominantly outcrossing domesticated *C. elegans* population.

The experimental populations were cultured following Teotonio et al. (2012). Population samples were revived from -80°C cryopreserved stocks containing at least 10^4^ individuals, and were expanded for two generations in a common environment. The populations were cultured in 9 cm Petri dishes containing NGM-lite agar (US Biological) with 25 mM NaCl and a 100 µL lawn of *E. coli* (HT115) as a food source and were kept at 20°C and 80% humidity. Populations underwent passaging with a 4-day non-overlapping life cycle from L1 larvae to mature adults. During experimental evolution, NaCl concentration was increased to 230 mM to create a challenging environment for populations to adapt to, referred to as the “high salt” environment (Parée et al., 2025). An increase in NaCl concentration reduces fertility and induces developmental delay (Theologidis et al., 2014). At each generation, synchronized starved L1 larvae were seeded in Petri dishes at a density of 10^3^ individuals per plate in 10 plates. L1 densities were estimated by counting the larvae in 15 µL of M9 under a Nikon SM1500. After 72 hours of growth and reproduction, adults and embryos were collected in M9 buffer (22 mM KH2PO4, 42 mM Na2HPO4, 85 mM NaCl, and 1 mM MgSO4), and subjected to a 5-minute treatment with 20 mM KOH:0.6% NaClO. This “bleach/hatch-off” protocol kills adults and larvae while allowing embryo survival. Following three washes in M9, the embryos were incubated in M9 at 20°C for 24 hours. L1 larvae are then seeded in fresh petri dishes for the next generation. Whenever needed, outcrossing was prevented by impairing males’ locomotion by adding 2 mL of M9 to the petri dishes 48h post-L1. This treatment forces hermaphrodites to self-fertilize, leading to a significant drop in male frequencies in the next generations. This operation could be repeated a few times during experimental evolution.

In total, 42 populations were experimentally evolved across two blocks Table S1. Note that the average selfing rates in the second block are lower but still overlap with the first block, allowing for modeling a block effect (see below).

### *rec-1* allele frequencies

*rec-1* allele frequencies were measured following the qPCR melting curve analysis protocol developed and presented in (Parée et al., 2025), which takes advantage of the distinct melting temperature of the two *rec-1* alleles to measure their proportion in pooled DNA of a population. Briefly, DNA was extracted and purified from whole-populations’ adult debris following the bleach/hatch-off protocol. *rec-1* alleles were amplified on a qPCR instrument, and amplicon were subsequently melted. The melting profile of each sample was then decomposed by an algorithm and compared to calibration samples, allowing for estimation of the *rec-1* allele frequency (*p*_*mutant*_) in the experimental sample.

### Data analysis

All analyses were performed in R (R Core Team, 2021), and figures were generated using the `ggplot2` package (Wickham and Sievert, 2009). The effect of the self-fertilization rate on *rec-1* mutant frequency trajectories was tested using a generalized linear mixed model (GLMM) implemented in the `glmmTMB` R package (Brooks et al., 2017):

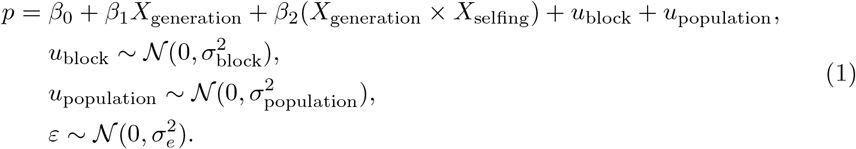

Where *p* is the *rec-1* mutant frequency, *β* are the fitted coefficients for the fixed continuous variables *X*, and *u* are the random effects. As we do not have count data but allele frequencies, we modeled the response using a beta distribution, which is appropriate for continuous proportions bounded between 0 and 1. *X*_selfing_ is a continuous variable corresponding to the mean selfing rate of populations, except for Figure 1D where it is a three-level categorical variable of arbitrary selfing classes (low, intermediate, high). P-values for model terms were obtained by likelihood ratio tests that compared a null model lacking the term. The term testing the effect of selfing-dependent selection of *rec-1* is *β*_2_, the effect of the interaction between selfing and the generation on *rec-1* frequencies. For selfing classes, estimated marginal means of selection coefficients and pairwise comparisons using the Scheffe method were implemented using the emmeans R package (Lenth, 2024).

Because selfing rates were unbalanced across experimental blocks and allele frequency trajectories tended to decline more strongly in the second block, we verified that the observed selfing-dependent selection was not driven by the block structure. To do so, we replaced the random block effect with a fixed interaction between block and generation:

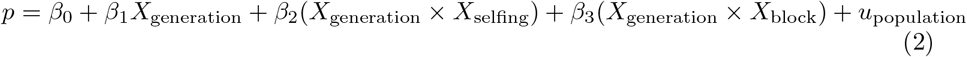

Selfing-dependent selection term (*β*_2_) is significant in this second model (P-value = 1.87 × 10^*−*2^). We also present, for each population, selection coefficients (*s*) obtained by solving the following binomial generalized linear model, implemented with the glm R function, set as *family=binomial* :

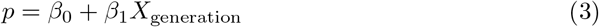

Where the selection coefficient is given by the fitted *β*_1_, and corresponds to the average rate of change of mutant frequency per generation on a *logit* scale. Because genetic drift is expected to increase with selfing (Nordborg and Donnelly, 1997), we tested the effect of the selfing rate on populations’ selection coefficient using the generalized least squares framework through the nlme::gls R function, which allows for explicit modeling of heteroskedasticity (Pinheiro, 2011).

### Simulations

Individual-based simulations of experimental evolution were run using SLiM 4.0.1 (Haller and Messer, 2023) using the scripts presented in Parée et al. (2025). We modeled an androdioecious diploid population of a size *N* = 10^3^ with hermaphrodites capable of both self-fertilization or outcrossing with males. In each generation, offspring genotypes were formed by combining two parental gametes. The first gamete was sampled from a hermaphrodite, with individuals weighted by their fitness. For self-fertilized offspring, the second gamete originated from the same hermaphrodite. For outcrossed offspring, the second gamete was sampled from a male, again weighted by fitness. Gametes were generated by recombining the two haploid genomes of each parent. Each homologous chromosome pair had a 50% chance of undergoing a single crossover event, corresponding to a genetic map length of 50 cM per chromosome. Depending of the allele identity at the recessive modifier locus, recombination break-points were sampled along the chromosome with probabilities corresponding on the linkage maps of wild-type or mutant *rec-1* alleles (Parée et al., 2024).

Haplotypes found in the experimental populations (Noble et al., 2021) were simulated for a subset of 1,000 randomly sampled SNV equally distributed on chromosomes I, II, III. Selection coefficients were randomly assigned to one of the alleles at each locus by sampling from a normal distribution centered at zero. A broad range of standard deviations was tested to identify the value yielding a genetic variance in fitness of 0.045 in the initial population. Individual fitness (*W*) in these simulated ancestral populations was modeled under a multiplicative selection with codominance:

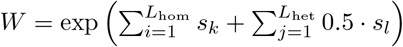

where *s* denotes the selection coefficient at the *k*^th^ homozygous or *l*^th^ heterozygous locus, among *L*_hom_ and *L*_het_ loci, respectively. Before simulations, negative LD, expected to be already present in these populations because of past selection, was generated through a burn-in of 100 generations under the wild-type recombination maps and negative epistasis (intermediate genotypes have higher fitness, extreme genotypes have lower fitness).

## Data and archiving

Data, and R code for analysis will be available upon publication.

## Acknowledgments

We thank H. Gendrot and V. Pereira for help with nematode handling, and L. Noble and F. Mallard for help with the SLiM simulations. We also thank C. Haag, T. Lenormand, F. Mallard, and M. Rockman for discussion.

## Author Contributions

TP: Conceptualization, Data curation, Investigation, Software, Formal analysis, Writing; NSC: Data curation, Investigation, Validation; DR: Investigation, Funding acquisition, Writing. HT: Conceptualization, Resources, Supervision, Funding acquisition, Investigation, Writing, Project administration.

## Supplementary

**Fig S1.**
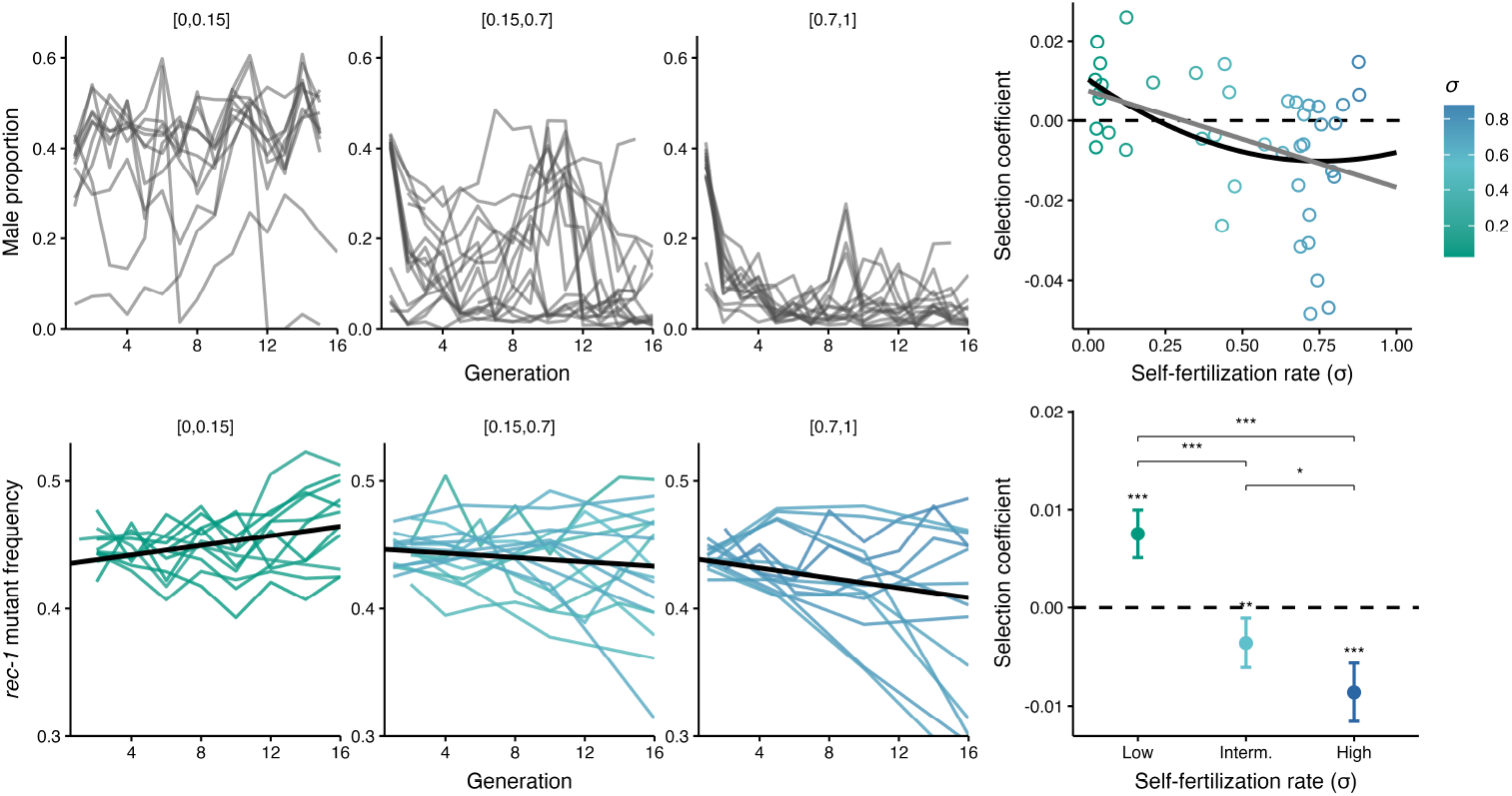
Experimental evolution with selfing binned by its harmonic mean. *A*. Male proportion during experimental evolution. Each line is an experimental population. The populations were classified into three groups based on the harmonic mean of their self-fertilization during evolution. *rec-1* allele frequency during experimental evolution. Each line is a replicate population. *C*. Each dot is the observed selection coefficient of the *rec-1* mutant for a given population. Lines represent the fitted regression line from the linear model with (black) and without (grey) a quadratic term. *D*. Dots and error bars are the estimated marginal mean and 95% confidence interval for each selfing rate bin estimated from a linear mixed model.

**Table S1.** Experimental populations. Asterisks indicates populations already presented in (Parée et al., 2025). Note that the gap between SMR18 and SMR50 is because those are chronological code internal to the lab and an unrelated experiment took place between the two blocks.

| Population | Experimental Block | Mean Selfing Rate ($\sigma$) |
| --- | --- | --- |
| SMR1* | 1 | 0.29 |
| SMR2 | 1 | 0.82 |
| SMR3* | 1 | 0.13 |
| SMR4 | 1 | 0.74 |
| SMR5* | 1 | 0.12 |
| SMR6 | 1 | 0.69 |
| SMR7* | 1 | 0.11 |
| SMR8 | 1 | 0.67 |
| SMR9* | 1 | 0.10 |
| SMR10 | 1 | 0.89 |
| SMR11* | 1 | 0.14 |
| SMR12 | 1 | 0.89 |
| SMR13* | 1 | 0.17 |
| SMR14 | 1 | 0.85 |
| SMR15* | 1 | 0.11 |
| SMR16 | 1 | 0.76 |
| SMR17* | 1 | 0.15 |
| SMR18 | 1 | 0.55 |
| SMR50 | 2 | 0.51 |
| SMR51 | 2 | 0.48 |
| SMR52 | 2 | 0.56 |
| SMR53 | 2 | 0.57 |
| SMR54 | 2 | 0.53 |
| SMR55 | 2 | 0.51 |
| SMR60 | 2 | 0.84 |
| SMR61 | 2 | 0.84 |
| SMR62 | 2 | 0.86 |
| SMR63 | 2 | 0.81 |
| SMR64 | 2 | 0.88 |
| SMR65 | 2 | 0.83 |
| SMR66 | 2 | 0.84 |
| SMR67 | 2 | 0.84 |
| SMR68 | 2 | 0.86 |
| SMR69 | 2 | 0.83 |
| SMR70 | 2 | 0.82 |
| SMR71 | 2 | 0.87 |
| SMR72 | 2 | 0.81 |
| SMR73 | 2 | 0.83 |
| SMR74 | 2 | 0.82 |
| SMR75 | 2 | 0.82 |
| SMR76 | 2 | 0.83 |
| SMR77 | 2 | 0.84 |

